# A *Streptococcus* quorum sensing system enables suppression of innate immunity

**DOI:** 10.1101/2020.12.03.411157

**Authors:** Kate Rahbari, Jennifer Chang, Michael Federle

**Affiliations:** Department of Microbiology and Immunology; University of Illinois at Chicago, Chicago, IL, 60607; USA; Department of Pharmaceutical Sciences; University of Illinois at Chicago, Chicago, IL, 60607; USA

**Keywords:** Immunosuppression, innate immunity, macrophage, pheromone, biosynthetic gene cluster, NFkB, TLR, cytokines

## Abstract

Some bacterial pathogens utilize cell-cell communication systems, such as quorum sensing (QS), to coordinate genetic programs during host colonization and infection. The human-restricted pathosymbiont *Streptococcus pyogenes* (Group A Streptococcus, GAS) uses the Rgg2/Rgg3 QS system to modify the bacterial surface, enabling biofilm formation and lysozyme resistance. Here, we demonstrate that innate immune cell responses to GAS are substantially altered by the QS status of the bacteria. We found that macrophage activation, stimulated by multiple agonists and assessed by cytokine production and NFκB activity, was substantially suppressed upon interaction with QS-active GAS but not QS-inactive bacteria. Neither macrophage viability nor bacterial adherence were seen as different between QS activity states, yet TNFα, IL-6, and IFNβ levels and NFκB reporter activity were drastically lower following infection with QS-active GAS. Suppression required contact between viable bacteria and macrophages. A QS-regulated biosynthetic gene cluster (BGC) in the GAS genome, encoding several putative enzymes, was also required for macrophage modulation. Our findings suggest a model wherein upon contact with macrophages, QS-active GAS produce a BGC-derived factor capable of suppressing inflammatory responses. The suppressive capability of QS-active GAS is abolished after treatment with a specific QS inhibitor. These observations suggest that interfering with the ability of bacteria to collaborate via QS can serve as a strategy to counteract microbial efforts to manipulate host defenses.

**Importance:** *Streptococcus pyogenes* is restricted to human hosts and commonly causes superficial diseases such as pharyngitis; it can also cause severe and deadly manifestations including necrotizing skin disease or severe post-infectious sequelae like rheumatic heart disease. Understanding the complex mechanisms used by this pathogen to manipulate host defenses could aid in developing new therapeutics to treat infections. Here, we examine the impact of a bacterial cell-cell communication system, which is highly conserved across *S. pyogenes*, on host innate immune responses. We find that *S. pyogenes* uses this system to suppress macrophage pro-inflammatory cytokine responses. Interference with this communication system could serve as a strategy to disarm bacteria and maintain an effective immune response.

## Introduction

Bacteria possess eloquent mechanisms to subvert host recognition and overcome the multiple barriers of human immune defenses. Strategies for survival have evolved to incorporate complex signaling systems that regulate genetic programs necessary to adapt to multiple environments and threats encountered in the host. Cell-cell communication systems, such as quorum sensing (QS), allow bacteria to disseminate information about their environment, density, and metabolism through the transmission of extracellular signals. QS systems invoke social pressures that result in the synchronization of genetic programs across the population. This synchronization enables bacteria to function as a multicellular group better equipped to perform activities such as forming biofilms, producing virulence factors, or taking up foreign DNA.^1^

The bacterium *Streptococcus pyogenes* (Group A Strep, GAS) is a serious pathogen responsible for a high global burden of disease, including 616 million annual cases of GAS pharyngitis and 1.78 million annual cases of severe diseases like necrotizing fasciitis, toxic shock syndrome, rheumatic heart disease, and glomerulonephritis.^2^ GAS utilizes several QS systems, including RopB (Rgg1), Rgg2/3, ComR (Rgg4), and Sil. These systems have been shown to regulate virulence factors, biofilm production, competence genes, and invasive disease, respectively.^1^ Notably, the Rgg2/3 system is conserved across all serotypes of GAS, indicating its importance for GAS survival and making it an attractive system to study.

The transcriptional regulators Rgg2 and Rgg3 respond to two functionally equivalent short hydrophobic peptide ligands (SHP2 and SHP3, collectively referred to as SHP) in a concentration dependent manner. When concentrations of SHP are low, the repressive activity of Rgg3 predominates at target promoters. When SHP reaches a critical concentration (~1 nM in culture conditions), it binds to Rgg2 and Rgg3 causing *inactivation* of Rgg3 (leading to de-repression of target genes) and *activation* of Rgg2 (leading to transcriptional activation of target genes). Genetic tools have helped identify two genetic loci that are directly controlled by the Rgg2/3 system: (1) the region adjacent to and including *shp2*, including *spy49_0414c* (*stcA*) and (2) the region adjacent to and including *shp3* and *spy49_0450-0460*. Rgg2/3-mediated regulation of *stcA* promotes *aggregation* and *biofilm development* (two phenotypes that require adhesive surface structures) and *resistance to lysozyme* (an antimicrobial host factor that targets components of the bacterial cell wall).^3, 4^ Because the surface of GAS interfaces with host cells, we hypothesized that Rgg2/3-mediated alterations to GAS would result in altered host immune responses.

Early innate immune responses to GAS are critical for control of infection.^5^ Pharmacological and genetic depletion studies established the importance of macrophages, dendritic cells (DCs), and neutrophils in initiating the innate immune response against GAS via distinct pathways.^5–8^ Macrophages are particularly important, as *in vivo* depletion or blockage of phagocytosis both drastically enhance susceptibility of mice to GAS.^5, 6^ In response to GAS, macrophages upregulate expression of genes involved in inflammation, survival, and production of oxygen radicals, which help them eliminate GAS efficiently.^9^ Signaling through MyD88 in macrophages and dendritic cells is required for cytokine responses and GAS clearance, illustrating the importance of pathogen sensing through TLRs and IL1R in response to infection.^10–13^ Downstream of MyD88, a cascade of events leads to the translocation of NFκB family proteins to the nucleus, where they regulate expression of genes required to promote inflammation, including cytokines, adhesion molecules, and cell growth or death factors.

Because the Rgg2/3 QS system alters surface properties of GAS, this study aimed to determine the consequence on immune responses. Macrophages were infected with isogenic GAS mutants lacking either the transcriptional repressor, *rgg3* (QS-locked ON) or the transcriptional activator, *rgg2* (QS-locked OFF). Here, we report that Rgg2/3 activation limits pro-inflammatory cytokine responses to GAS. Moreover, QS activation resulted in suppression of cytokine responses induced by several TLR agonists. Expression of the putative biosynthetic gene cluster (BGC) downstream of *shp3* was found to be required for this phenotype. Overall this study has identified a new role for Rgg2/3 QS in manipulating innate immune responses to GAS.

## Results

### Macrophage responses to GAS are attenuated when Rgg2/3 QS is active

Macrophages respond to GAS in part by activating NFκB, a transcription factor capable of regulating expression of many pro-inflammatory cytokines. To determine if the GAS QS state impacts macrophage responses, pro-inflammatory responses were measured following *in vitro* infections with wild-type (WT), Δ*rgg3* (QS locked ON), or Δ*rgg2* (QS locked OFF) GAS. Infection with WT or Δ*rgg2* resulted in activation of NFκB and production of high amounts of TNFα and IL-6 (**Fig 1A-B**). Interestingly, infection with Δ*rgg3* led to approximately 4-fold decreased NFκB reporter activity and 20-fold decreased TNFα and 25-fold decreased IL-6 production compared to infection with WT or Δ*rgg2* (**Fig 1A-B**). To determine if decreased activity was due to differences in macrophage cytotoxicity, extracellular release of the cytoplasmic enzyme lactate dehydrogenase (LDH) was measured as an indicator of plasma membrane damage. LDH release was minimal in response to WT and mutant GAS strains after 8 hours of infection (**Fig 1C**). Additionally, infecting macrophages with increasing MOI of Δ*rgg2* resulted in dose-dependent activation of NFκB (**Fig 1D**). Increasing the MOI of Δ*rgg3*, however, failed to increase NFκB responses, and instead they remained attenuated even at the highest MOI tested.

**Figure 1.**
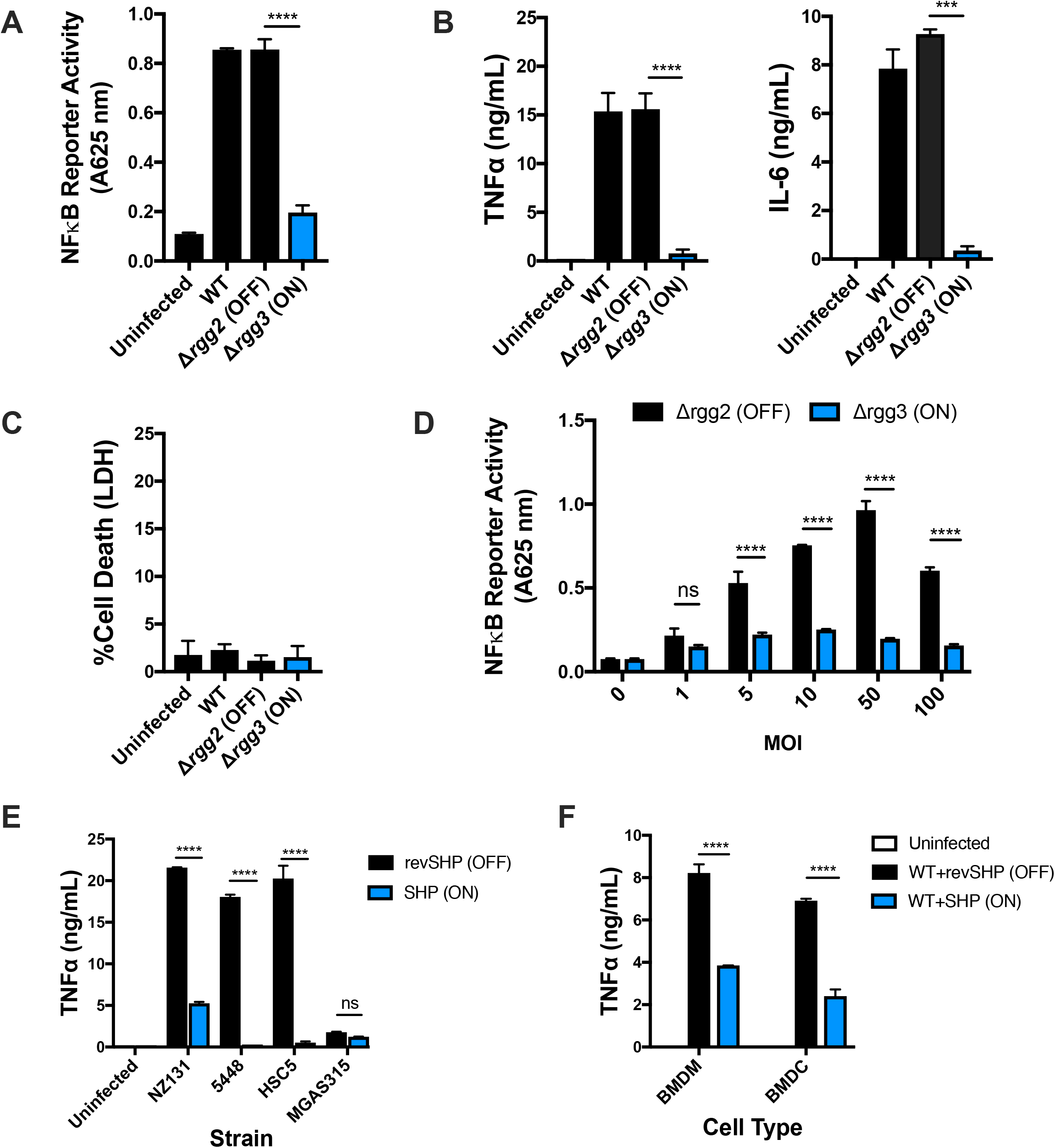
Macrophage responses to GAS are attenuated when Rgg2/3 QS is active. (**A**) NFκB responses after infecting RAW246.7 cells containing a chromosomally integrated NFκB-inducible secreted embryonic alkaline phosphatase reporter (RAW-Blue cells) with wild-type (WT) Δ*rgg2* (QS-OFF), or Δ*rgg3* (QS-ON) GAS. All cells were infected with MOI 10 unless otherwise indicated. Reporter activity (absorbance 625 nm) is shown. (**B**) TNFα and IL-6 production by RAW264.7 cells 8 hours after infection. (**C**) LDH release as a measurement of macrophage cell death 8 hours after infection. Percentage of dead cells was quantified using a 100% lysed cell contFκB activity after infecting RAW-Blue crol. (**D**) Nells for 20 hours with different multiplicities of infection (MOI) of Δ*rgg2* (black bars) or Δ*rgg3* (blue bars). (**E**) TNFα production by RAW264.7 cells 8 hours after infection with different serotypes of GAS grown in the presence of revSHP (black bars) or SHP (blue bars). (**F**) TNFα production by BMDM or BMDC 8 hours after infection with GAS (strain NZ131) grown in the presence of revSHP (black bars) or SHP (blue bars). Means ± SD are shown from a representative of three independent experiments conducted in triplicate. ***p=0.001; ****p<0.0001 by two-tailed unpaired t test (A-B) or ordinary one-way ANOVA with Tukey’s multiple comparisons test (C-E).

The chemically defined medium (CDM) used to grow the bacteria in the laboratory lacks SHP pheromones and other signals capable of activating Rgg2/3 QS; thus the wild-type strain is QS-OFF when cultured in this medium.^3, 14^ We have previously shown that exogenous addition of synthetic SHP pheromone during growth of WT culture activates Rgg2/3 QS within minutes.^3^ Macrophage inoculation with SHP-activated WT GAS (strain NZ131) led to attenuated TNFα production similarly to that of the genetically QS locked ON mutant, Δ*rgg3* (**Fig 1E**). Macrophage responses WT GAS grown in the presence of the inactive reverse sequence of SHP (revSHP) mimicked that of the genetically QS locked OFF mutant, Δ*rgg2*.

Rgg2/3 QS is conserved across all sequenced serotypes of GAS. GAS serotypes are classified based on differences in M protein, a surface protein and major virulence factor. The strain NZ131 (serotype M49) was used for a majority of this work because it is a highly transformable strain and excellent tool for genetic manipulation of GAS. To confirm that the immunomodulatory phenotype is not restricted to NZ131, macrophage TNFα responses were measured following infection with other wild-type strains: HSC5 (M14), 5448 (M1), and MGAS315 (M3). These particular strains were chosen as representatives of diverse M types. Furthermore, we previously confirmed that Rgg2/3 activation leads to cell surface changes (lysozyme resistance and biofilm formation) in both HSC5 and MGAS315.^14^ Strain 5448 was added as a representative M1T1 globally disseminated, clinically relevant strain. With the exception of MGAS315, activation of Rgg2/3 QS by the addition of exogenous SHP resulted in reduced macrophage TNFα production in response to all other strains tested (**Fig 1E**). Curiously, minimal TNFα was produced in response to MGAS315, even without the activation of QS. This may be due to the strain’s high expression of capsule, a virulence factor that prevents adherence and phagocytosis.^15^

To determine if the differential cytokine response is restricted to RAW264.7 macrophages, the effect of QS activation on cytokine production was also examined following *in vitro* infection of primary mouse bone marrow-derived macrophages (BMDM) and dendritic cells (BMDC). TNFα production by BMDM and BMDC was attenuated in response to Δ*rgg3*, consistent with responses seen in RAW264.7 cells (**Fig 1F**). These data suggest that Δ*rgg3* is able to manipulate cytokine production in multiple innate immune cell types, indicating the potential relevance and applicability of interfering with QS as therapeutic strategy.

### QS-ON GAS actively suppresses macrophage inflammatory responses

To determine if QS enabled GAS to evade detection (i.e. hide) or rather to actively downregulate (i.e. suppress) inflammatory responses, macrophages were infected with mixed populations of Δ*rgg3* and Δ*rgg2*, and NFκB activity or cytokine production was measured. Following mixed infection with both Δ*rgg3* and Δ*rgg2*, NFκB activity remained low and was similar to that of Δ*rgg3* single infection (**Fig 2A**). Decreasing the MOI of Δ*rgg3* in mixed infections led to a dose-dependent increase in production of all cytokines tested (TNFα, IL-6, IFNβ). Surprisingly, cytokine production remained attenuated even when Δ*rgg3* was outnumbered 5 to 1 by Δ*rgg2* (**Fig 2B**). TNFα production by primary mouse BMDM and BMDC was also attenuated following mixed infections with equal MOI of Δ*rgg2* and either Δ*rgg3* or WT grown in the presence of SHP (**Fig 2C**). Together these data suggest that QS enables GAS to suppress macrophage cytokine responses.

**Figure 2.**
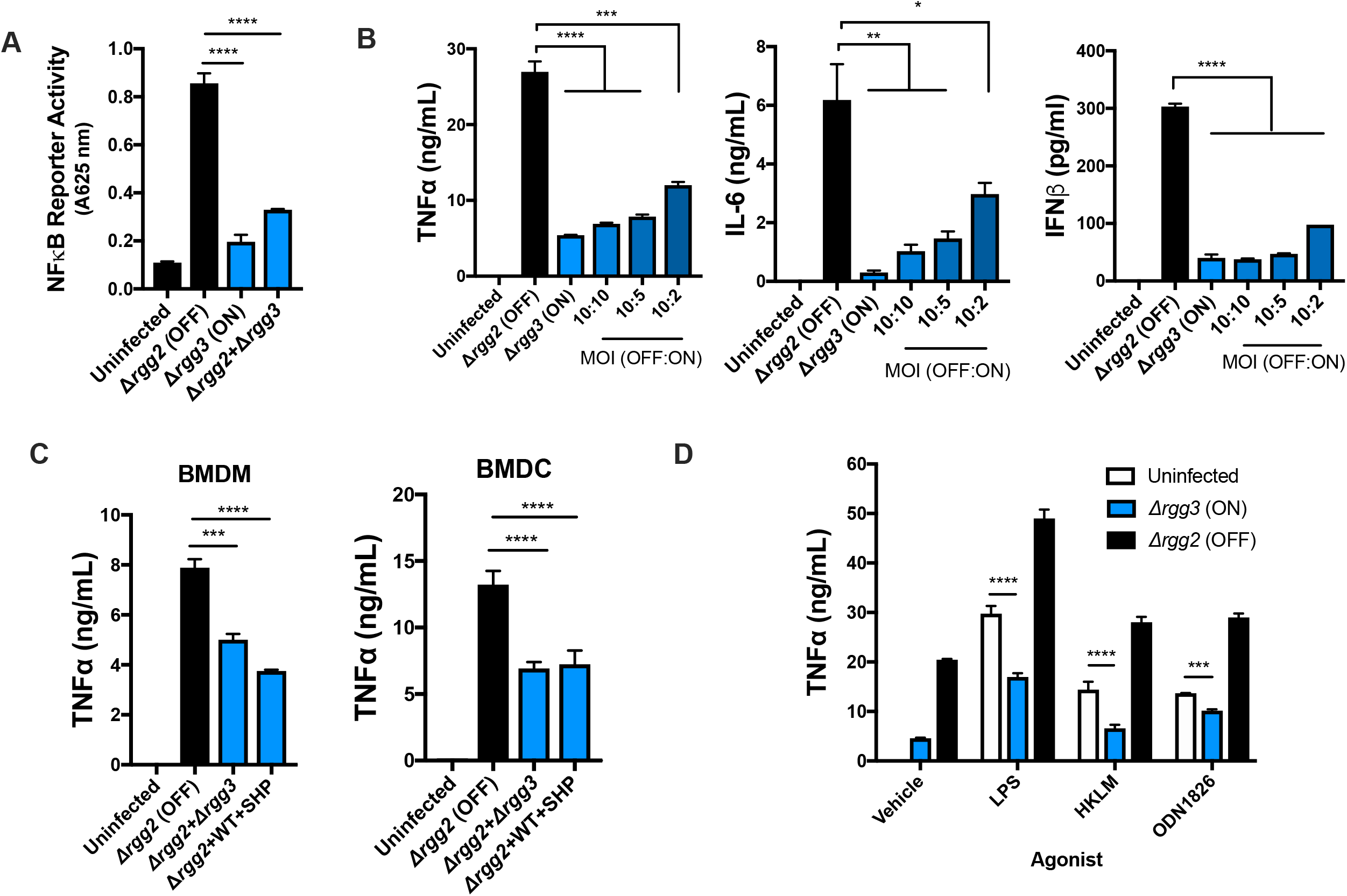
QS-ON GAS actively suppresses inflammatory responses. (**A**) NFκB activity after infecting RAW-Blue cells with Δ*rgg2*, Δ*rgg3*, or both. (**B**) TNFα, IL-6, and IFNβ production after RAW264.7 cells were infected with the described MOIs of Δ*rgg2* and Δ*rgg3*. (**C**) TNFα production by BMDM or BMDC 8 hours after infection with MOI 10 of Δ*rgg2* or co-infected with MOI 10 of each Δ*rgg2* and Δ*rgg3* or Δ*rgg2* and WT grown in the presence of SHP. (**D**) TNFα production after RAW264.7 cells were co-incubated with TLR agonists and Δ*rgg3* or Δ*rgg2*. LPS, lipopolysaccharide (TLR4); HKLM, Heat killed *Listeria monocytogenes* (TLR2); ODN1826, CpG oligodeoxynucleotide (TLR9). Means ± SD are shown from a representative of two (D) or three (A-C) independent experiments conducted in triplicate. *p < 0.05; **p < 0.005; ***p < 0.001; ****p < 0.0001, by ordinary one-way ANOVA with Tukey’s multiple comparisons test (A-C) or two-way ANOVA with Sidak’s multiple comparisons test (D).

The specific pathogen-associated molecular patterns (PAMPs) responsible for inducing pro-inflammatory responses to GAS are incompletely described; however, it has been well established that both the MyD88-NFκB pathway and the type I IFN pathway are required for clearance.^11, 13^ We wondered whether QS would enable GAS to suppress stimulation by TLR agonists. TLR agonist-stimulated macrophages produced significantly higher amounts of TNFα when applied simultaneously with Δ*rgg2* (**Fig 2D**). However, agonist treatment together with Δ*rgg3* infection resulted in decreased TNFα production in response to lipopolysaccharide (LPS; TLR4), heat-killed *L. monocytogenes* (HKLM; TLR2), and CpG oligodeoxynucleotide (ODN1826; TLR9). Together, these data suggest that GAS suppresses NFκB-mediated pro-inflammatory responses downstream of TLR stimulation.

### Suppression requires contact with live QS-ON GAS

To localize the factor(s) responsible for QS-regulated cytokine modulation, the ability of secreted/extracellular components and surface-associated components to alter NFκB activity was tested. In other pathogenic bacteria, QS molecules themselves have been shown to possess immunomodulatory properties toward host cells. For example, treatment of epithelial cells with QS molecules produced by *Pseudomonas aeruginosa* (i.e. homoserine lactones, quinolones, and phenazines) results in differential regulation of cytokine expression via sensing by the host receptor AhR.^16^ Macrophage treatment with the Rgg2/3 QS molecule SHP failed to increase NFκB activity compared to untreated controls. Therefore, we tested whether addition of SHP to Δ*rgg2* infected macrophages was sufficient to attenuate NFκB activity. We saw that addition of 200 nM of either SHP or revSHP to macrophages was unable to suppress NFκB activity (**Fig 3A**). These data suggest that although SHP signals induce Rgg2/3 signaling, they are not directly responsible for the observed immunomodulatory effect.

**Figure 3.**
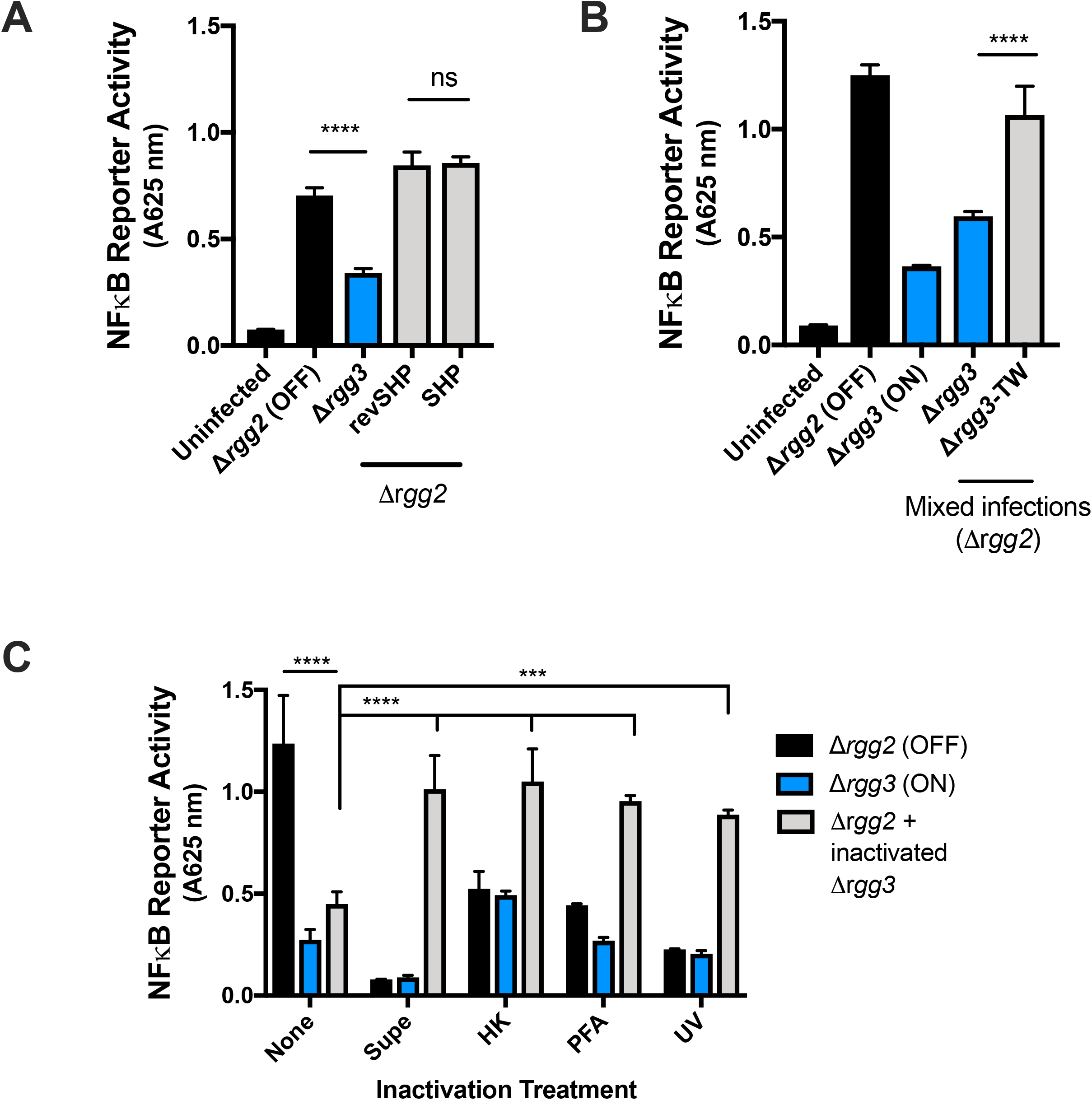
Suppression requires contact with live QS-ON GAS. (**A**) NFκB activity after stimulating RAW-Blue cells with MOI 10 Δ*rgg2* and treating with 200 nM SHP or revSHP, or MOI 10 Δ*rgg3*. (**B**) NFκB activity after infecting RAW-Blue cells directly with the described strains, or with Δ*rgg3* separated by a trans-well with 0.4 um pores (TW). (**C**) NFκB activity after infecting RAW-Blue cells with live Δ*rgg2* and/or GAS that was inactivated with heat treatment (HK), short wave ultraviolet light treatment (UV), or 4% paraformaldehyde (PFA), or OD-normalized sterile filtered supernatants (supe). Means ± SD are shown from a representative of three independent experiments conducted in triplicate. ****p < 0.0001 by two-way ANOVA with Sidak’s multiple comparisons test (A-B) or ****p < 0.0001, by ordinary one-way ANOVA with Tukey’s multiple comparisons test (C-D).

To determine whether other secreted factors could enable QS-ON GAS to suppress NFκB activity, macrophages were directly stimulated with Δ*rgg2* GAS, and Δ*rgg3* GAS was added to macrophages either directly or separated by a 0.4 μm pore trans-well membrane (TW). This size pore prohibited Δ*rgg3* GAS from directly contacting macrophages, and it simultaneously allowed secreted factors to pass through the membrane. We determined that when Δ*rgg3* GAS was added above the TW (Δ*rgg3-*TW), it was unable to suppress NFκB activity (**Fig 3B**, white bar). This indicated that macrophage suppression is a contact-dependent phenomenon. Furthermore, sterile filtered supernatants from Δ*rgg3* cultures were added to macrophages infected with Δ*rgg2* and failed to suppress NFκB activity, further indicating that GAS culture supernatants lack the factor responsible for suppressing macrophage responses (**Fig 3C**).

Our previous work illustrated that cell surface properties of GAS are altered via Rgg2/3 QS. ^3, 4, 14^ To examine if these surface changes include alterations of the repertoire or display of antigens, macrophages were infected with GAS inactivated by different methods, including heat (heat killed, HK), short wave ultraviolet light (UV), and 4% paraformaldehyde (PFA) treatment. While heat has been shown to alter cell wall peptidoglycans and denature proteins, UV treatment damages DNA and RNA and is less likely to interfere with cell surface properties. PFA treatment crosslinks amines in proteins and other structures within the cell. We posited that if QS-induced surface changes are responsible for altered macrophage responses, infection with inactivated GAS would similarly result in differential inflammatory responses. After each inactivation treatment, bacteria were plated for enumeration to confirm loss of viability, and in each case >99.9% of cells were non-recoverable. Macrophages infected with inactivated GAS had lower NFκB activity compared to those infected with live GAS, and this was consistent across all treatments (**Fig 3C**). Inactivation of GAS with heat and UV eliminated the QS-dependent differences in NFκB activity (**Fig 3C**). Furthermore, inactivation of Δ*rgg3* by any treatment eliminated its ability to attenuate NFκB activity when added to cells stimulated with live, metabolically active Δ*rgg2* (**Fig 3C)**. This suggests that alterations in surface properties are unlikely responsible for the differences in macrophage responses to Δ*rgg3* GAS, though it is possible that these inactivation methods may have interfered with the unknown ligand responsible for immunosuppression. Cumulatively, these data indicate that immunomodulation by QS-ON GAS requires live bacteria and direct contact between the pathogen and host cells.

### Suppression does not require bacterial internalization

GAS initially interacts with macrophages through bacterial adherence and internalization. Though GAS was first thought to be an exclusively extracellular pathogen, much evidence has illustrated that it can also invade and even replicate intracellularly.^17, 18^ While it appears that contact is required for QS-mediated immunosuppression (**Fig 3B**), we next wondered whether GAS must also be internalized to suppress macrophage responses. Inhibition of macrophage actin polymerization by treatment with Cytochalasin D (CytD) is well described to prevent phagocytosis of bacteria, including *S. pyogenes*.^19^ To determine whether QS-mediated suppression requires internalization of GAS, macrophages were pre-treated with CytD and then infected with Δ*rgg3*. After killing extracellular GAS with gentamicin for 30 minutes and washing away any remaining GAS, antibiotic, and CytD, LPS or Δ*rgg2* was added to stimulate the cells. We showed that inhibiting phagocytosis had no effect on the ability of QS-ON GAS to suppress stimulation by LPS or Δ*rgg2* GAS (**Fig 4A**). Next, to determine if QS activity modulates the degree to which GAS adheres to macrophages, GAS was plated for CFU enumeration after infection and compared to the initial inoculum. As a negative control, the Δ*covR* mutant strain of GAS was also tested because this strain is well defined to overexpress capsule (an antiphagocytic and anti-adherence factor). Adherence to macrophages after 30 minutes of infection was not affected by the QS-state of GAS (**Fig 4B**).

**Figure 4.**
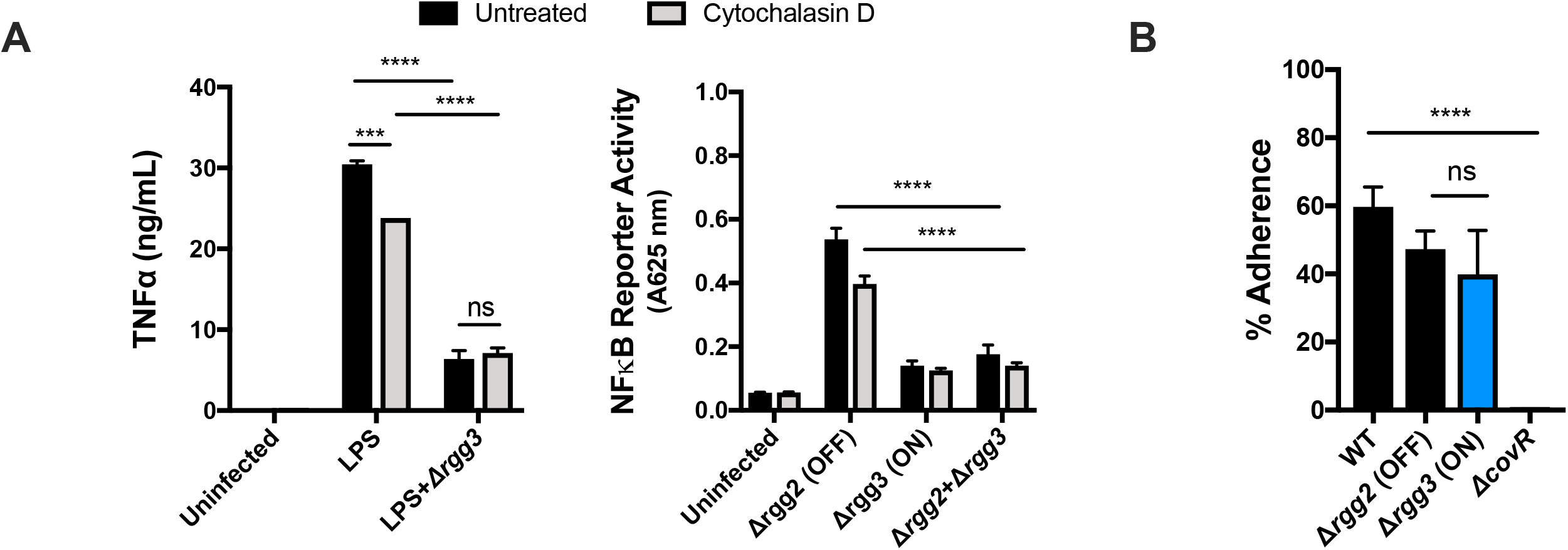
Suppression does not require bacterial internalization. (**A**) TNFα production (Left graph) or NFκB activity (Right graph) after RAW264.7 cells were pre-treated with Cytochalasin D and infected with Δ*rgg3* for 30 minutes, followed by stimulation with LPS or Δ*rgg2*. (**B**) GAS adherence to RAW264.7 cells after 30 minutes of infection. Adherence was calculated as the percentage of viable bacterial cells recovered compared to infection inoculum. Means ± SD are shown from a representative of two independent experiments conducted in triplicate. ***p=0.002, ****p < 0.0001, by two-way ANOVA with Sidak’s multiple comparisons-test (A) and ordinary one-way ANOVA with Tukey’s multiple comparisons test (B).

### A QS-regulated operon is required for cytokine suppression

Several virulence factors have been described as important in manipulating immune responses to GAS, including streptolysin O (SLO), *S. pyogenes* cell envelope protease (SpyCEP), M protein, and capsule, all of which interact with the bacterial surface.^20, 21^ To test whether any of these are involved, we activated the Rgg2/3 system by addition of SHP pheromone in isogenic strains lacking SLO (Δ*slo*), capsule (Δ*hasAB*), M protein (Δ*emm*), or SpyCEP (Δ*spycep*). When QS was activated, each of these strains maintained their capacity to suppress macrophage NFκB upon infection suggesting that these factors are not involved in the QS-dependent phenotype **(Fig 5A).**

**Figure 5.**
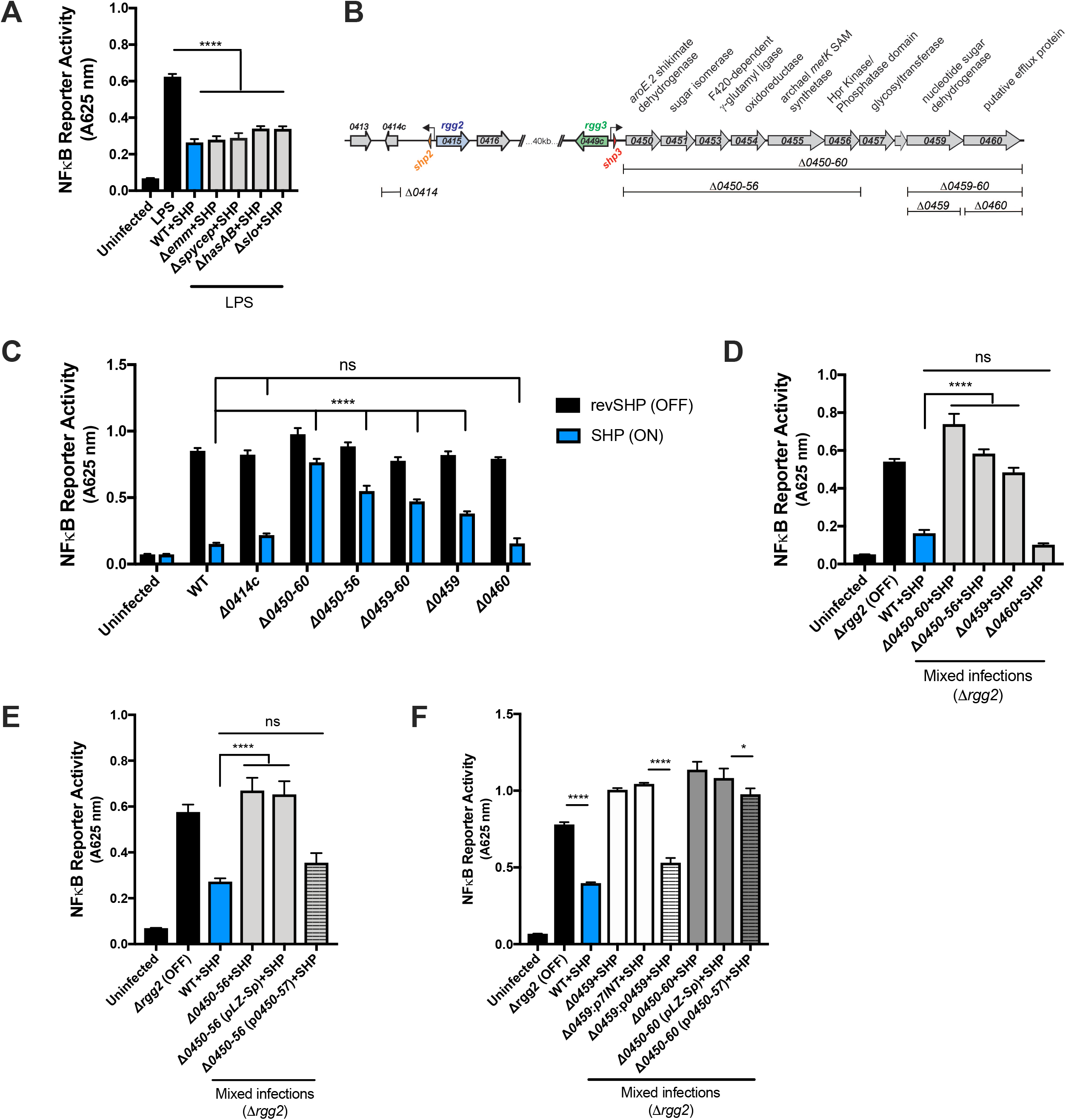
A QS-regulated biosynthetic gene cluster (BGC) is required for cytokine suppression. (**A**) NFκB activity of RAW-Blue cells infected with MOI 10 of GAS isogenic mutants lacking various virulence factor genes and grown in the presence of SHP. Following 30 minutes of infection, macrophages were stimulated with LPS. (**B**) Diagram depicting the QS-regulated genetic programs in GAS (strain NZ131), putative functions of the genes, and the isogenic mutants used in subsequent assays. (**C**) NFκB activity after RAW-Blue cells were infected with MOI 10 of the described GAS genetic mutant strains grown in the presence of either inactive reverse SHP (revSHP) or active SHP. (**D**) NFκB activity after RAW-Blue cells were co-infected with MOI 10 Δ*rgg2* and MOI 10 of the described BGC mutants grown in the presence of SHP. (**E-F**) NFκB activity after RAW-Blue cells were stimulated with MOI 10 Δ*rgg2* (**E**) or 100ng/ml LPS (**F**) and infected with BGC mutants and complementation strains. All GAS strains were grown in the presence of SHP to induce Rgg2/3 QS activity. Means ± SD are shown from a representative of three independent experiments conducted in triplicate. ****p < 0.0001, by ordinary one-way ANOVA with Tukey’s multiple comparisons test (A, D-F) or two-way ANOVA with Sidak’s multiple comparisons-test (C).

**Figure 6.**
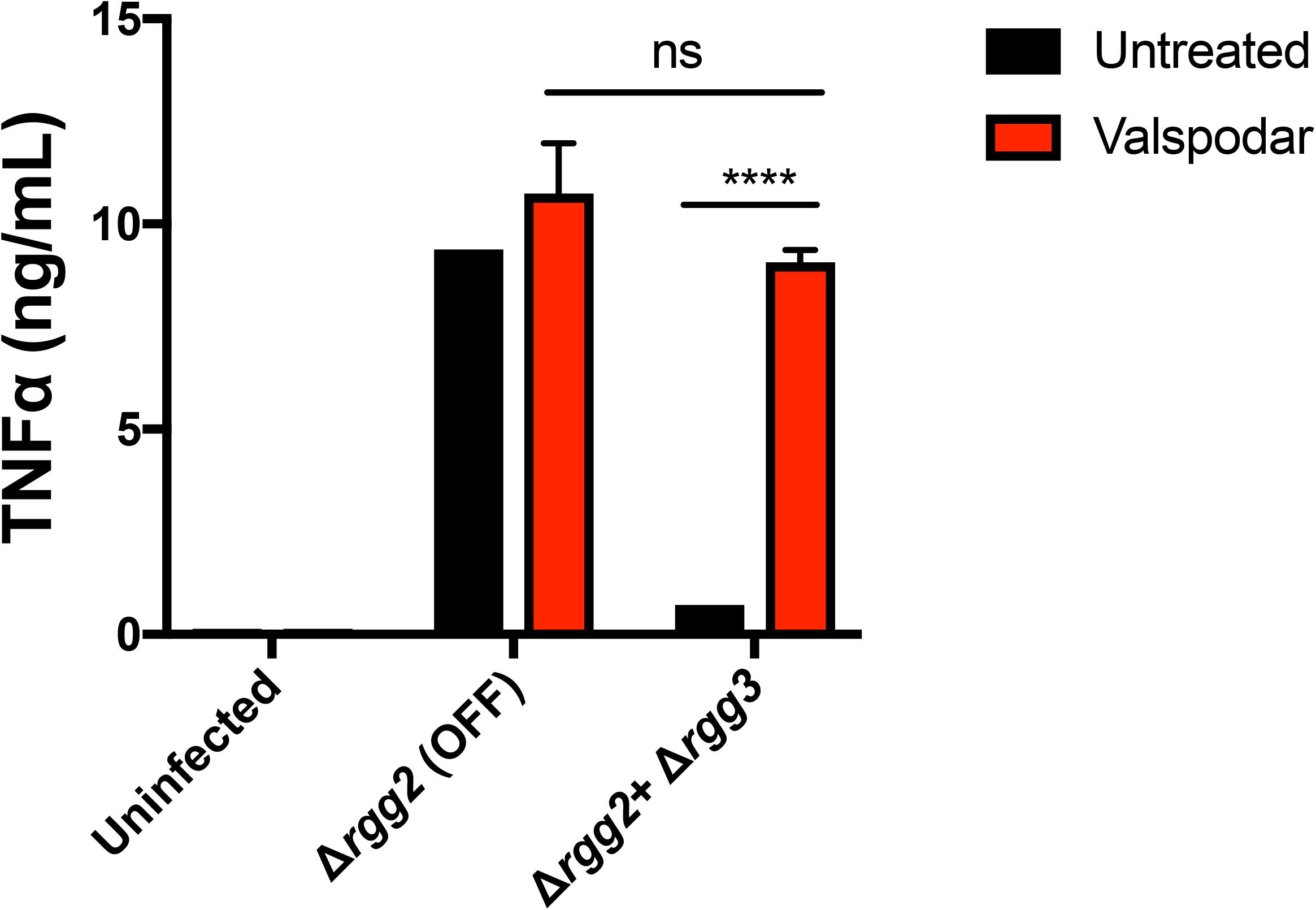
Rgg2/3 QS can be targeted pharmacologically to restore immunity. TNFα production after RAW264.7 cells were infected with GAS that was treated with or without the QS inhibitor valspodar (10 μM). Means ± SD are shown from a representative of three independent experiments conducted in triplicate. ****p < 0.0001, by two-way ANOVA with Tukey’s multiple comparisons-test.

The major genetic targets of the Rgg2/3 QS pathway include: (1*) spy49_0414c*, encoding the protein StcA and (2) *spy49_0450-0460,* encoding a putative biosynthetic gene cluster (BGC) (**Fig 5B**).^3^ We previously described that activation of Rgg2/3 QS mediates production of biofilms and resistance to the host antimicrobial factor lysozyme, two phenotypes that rely on surface modifications and were shown to require QS-regulation of StcA.^3, 4^ We wondered whether regulation of StcA also enabled suppression of inflammatory pathways in innate immune cells. Infecting macrophages with Δ*0414c and* Δ*0450-0460* grown in the presence of SHP (QS-ON) or inactive revSHP (QS-OFF) showed that while *spy49_0414c* is not required for QS-regulated attenuation of cytokine responses, *spy49_0450-0460* is required (**Fig 5C**). Further interrogation using isogenic mutants of various genes within this pathway determined that QS-ON GAS lacking either the first six genes (*spy49_0450-0456*) or the second to last gene, *spy49_0459*, failed to attenuate NFκB activity following single infections (**Fig 5C**). Interestingly, infection with QS-ON GAS lacking only *spy49_0460* attenuated NFκB activity to similar levels as infection with the WT control (**Fig 5C**). Consistent with this observation, mixed infections with QS-OFF GAS and WT or BGC mutants grown in the presence of SHP pheromone (QS-ON), demonstrated that BGC mutants significantly restored the NFκB response, suggesting these genes are involved in suppression (**Fig 5D**). A mutant strain lacking only *spy49_0460,* however, retained its ability to suppress NFkB activity, again suggesting this gene is not required for this activity.

Complementation of *spy49_0450-0457* or *spy49_0459* in the corresponding deletion strains was able to restore suppression capability (**Fig 5E-F**). However, complementation of *spy49_0450-0457* in Δ*0450-0460* failed to fully complement suppression, suggesting the importance of later genes in the BGC, such as *spy49_0459* (**Fig 5F**). Together, these data suggest that the previously described surface alterations mediated by *stcA* are not responsible for macrophage suppression by QS-ON GAS, and instead genes in the BGC are involved.

### Rgg2/3 QS can be targeted pharmacologically to restore immunity

The ability of QS-activated GAS to suppress host cytokine responses could be a new potential mechanism to target *in vivo* as an anti-virulence treatment strategy. Previous screening of a library of FDA approved drugs identified cyclosporine-A (CsA) as an inhibitor of Rgg2/3 QS signaling. The CsA analog valspodar, which lacks immunosuppressive effects, was also found to inhibit signaling.^22^ To determine if pharmacologically inhibiting QS could prevent immunosuppression, Δ*rgg3* and Δ*rgg2* cultures were treated with 10 μM valspodar and then applied to macrophages. Compared with untreated Δ*rgg3*, valspodar-treated GAS was no longer able to suppress cytokine production in macrophages. These data indicate that blocking QS-signaling can restore macrophage responses to GAS and eliminate their ability to suppress cytokine production.

## Discussion

This study presents a novel phenomenon whereby GAS utilizes an intercellular regulatory system to actively suppress NFκB activity and pro-inflammatory cytokine production in macrophages and dendritic cells. Bacteria incapable of responding to pheromone signals (QS-OFF mutants) lacked the ability to suppress immune cell responses. QS-ON GAS that were inactivated by heat treatment, fixation, or UV irradiation also lost their ability to suppress inflammatory responses. This indicates that suppression is an active process occurring at the interface of GAS and macrophages rather than presentation of a stable surface moiety by GAS. Cell-free culture supernatants also failed to suppress inflammatory responses, further supporting the notion that a secreted factor is not released into the surrounding milieu. Separation of bacteria from stimulated macrophages by a trans-well membrane also abolished suppression capacity of QS-ON GAS. Together, these data support a model whereby GAS contacts innate immune cells and, when QS signaling is active, presents or delivers a factor that interrupts host signaling pathways.

The Rgg2/3 QS system directly regulates two operons, each encoding a *shp* gene and additional coding sequences: *spy49_0414c* (*stcA*) and *spy49_0450-0460*, a putative biosynthetic gene cluster. Here, we were able to link immune cell suppression activity to expression of *spy49_0450-0460;* deletion of the operon abolished suppression. This locus is conserved across all sequenced GAS genomes and is comprised of 10 genes co-expressed from the *shp3* promoter. They encode various enzymes involved in amino acid biosynthesis, sugar metabolism, and cell wall biogenesis (**Fig 5B**). The operon was found to be repressed by the metal dependent regulator MtsR, and some of the genes were identified in screens for those involved in invasive infection and survival in blood.^23–25^ Yet, little remains known about this putative biosynthetic gene cluster, and similar gene sets are seen in only a few other instances in *S. porcinus*, *S. pseudoporcinus*, *Salinispora tropica*, and *Bacillus thuringiensis.*

Curiously, in *B thuringiensis*, the homologous gene cluster is implicated in the biosynthesis of a secreted toxin called thuringensin, a nucleoside analog predicted to have cytotoxic activity.^26^ We observed no cytotoxic impact to macrophages incubated with QS-ON or QS-OFF cells (**Fig 1C**); however, we hypothesize that the product of the *spy49_0450-0460* gene cluster could be a bioactive molecule produced in response to and directed at macrophages. The last gene of the operon, *spy49_0460* (*mefE*), encodes a putative efflux protein. The presence of this gene in the operon is consistent with the notion that it aids in delivery of a molecule. Surprisingly, however, deletion of *mefE* did not affect QS-mediated cytokine suppression in macrophages, refuting its potential involvement in the observed activity, unless a redundant pathway is capable of releasing a factor.

Bacteria use diverse strategies to modulate host responses. We hypothesize that GAS produces a QS-regulated factor that can manipulate host signaling pathways and subsequently shape cytokine responses. Upon TLR activation, a cascade of phosphorylation and ubiquitination events leads to degradation of the inhibitor IκB, allowing NFκB translocation to the nucleus where it regulates transcription. Here, we demonstrate that GAS suppresses stimulation at a step downstream of TLR activation and upstream of NFκB transcriptional induction. Several examples of NFκB-pathway disruptors have been described in Gram-negative pathogens, and include *Yersinia* YopJ, *E. coli* Nle proteins, and *Shigella* Osp proteins.^27^ Fewer examples have been described in Gram-positive pathogens which largely lack specialized secretion systems used to deliver factors directly to host cells. We found that QS-induced suppression of macrophages required contact with live GAS and was not found in supernatants, (**Fig 3**) This suggests the factor is not released from bacteria into extracellular spaces, but instead it might be delivered by a mechanism allowing translocation across the host membrane, such as that seen in Type III secretion systems (T3SS). In GAS, delivery of the virulence factor NAD-glycohydrolase (NADase) has been shown to occur in a mechanism functionally equivalent to T3SS, but requiring the pore-forming toxin, streptolysin O (SLO) for delivery.^28, 29^ We induced the Rgg2/3 system in a strain lacking SLO (Δ*slo*) and found it maintained its capacity to suppress NFκB **(Fig 5A)**. Thus, if Rgg2/3-dependent immune suppression involves a translocated factor, effector delivery may involve as-of-yet characterized delivery mechanisms.

The ability of *Staphylococcus aureus*, another Gram-positive pathogen, to actively interfere with host signaling components does not necessitate translocation of factors into host cells. For example, staphylococcal superantigen-like protein (SSL3) binds TLR2 and blocks ligand binding and TLR heterodimerization, thereby inhibiting macrophage stimulation.^30, 31^ Another protein, TirS, blocks TLR-dependent NFκB activation by mimicking and interfering with host TIR-containing adaptor proteins (like MyD88), and presumably must enter the host cell cytoplasm in order to function.^32^ How TirS gains access to the host cytosol is unresolved.

Although many GAS virulence factors facilitate immune evasion by various mechanisms, none have been described to suppress NFκB responses. Multiple virulence factors in GAS can suppress cytokines post-translationally. For example, SLO induces host degradation of IL-1β via ubiquitination, and SpyCEP directly degrades IL-8 via proteolytic cleavage.^33, 34^ Here, we describe a distinct mechanism of suppression upstream of these pathways. Interestingly, the strain MGAS315 was the only strain tested which failed to suppress macrophage cytokine responses (**Fig 1E**). Further exploration of the differences between the MGAS315 strain and the others could help shed light on the bacterial factors responsible for the differences in macrophage responses. We suspect that differences in capsule, which prevents adherence to host cells, are involved because direct contact was required for suppression (**Fig 3**). We plan to examine this by deleting capsule genes in this strain and assessing its immunomodulatory potential. Interestingly, deleting capsule genes in NZ131 did not improve suppression compared to WT, suggesting there may be redundant factors involved.

Rgg2/3 QS is activated in response to environmental conditions found in host mucosal environments, including metal depletion and mannose utilization.^14^ It remains unclear what benefits are afforded by the additional layer of intercellular communication signaling. We recently demonstrated the importance of Rgg2/3 QS activation *in vivo* for nasopharyngeal colonization.^35^ Whether Rgg2/3-mediated immunosuppression was a contributing factor in the observed increased colonization rates of *rgg3* mutants (QS-ON) was not a component of these studies, but blocking NFκB signaling likely diminishes the efficacy of host strategies for bacteria eradication, including production of antimicrobial factors (lysozyme, cationic peptides), induction of autophagic responses, and recruitment of additional phagocytic and cytotoxic immune cells. Incorporating cell-cell signaling under threatening conditions likely reinforces bacterial compliance to engage defense mechanisms that improve probabilities of survival if immune responses are blunted.

As we gain deeper understanding of how QS systems are used by bacteria to enhance fitness attributes in the context of host colonization and/or infection, the more promising it becomes to target signaling activities pharmacologically as an alternative or supplementary therapeutic approach to antibiotics. We previously determined that the FDA-approved drugs cyclosporin A and valspodar have specific inhibitory activities against Rgg2/3 QS.^22^ We are encouraged by results shown here that blocking QS-activity with valspodar abolished its cytokine attenuation effects and restored macrophage inflammatory responses to GAS *in vitro*. Testing methodologies that incorporate anti-QS therapeutics remains a compelling strategy that could bolster the host’s intended immune response and diminish infections by this global pathogen.

## Acknowledgments

We thank Drs. Reid Wilkening and Juan Jimenez for strain construction, Dr. Benjamin Gantner for RAW264.7 cells, and Drs. Donna Macduff, Francis Alonzo, and David Ucker for helpful discussions. This study was supported by NIH R01AI091779 to M.J.F. and NIH F31AI147429 to K.R.

## Author Contributions

Conceptualization, data analysis, and funding acquisition, K.R. and M.J.F.; Investigation, K.R. and J.C.; Writing, K.R.; Reviewing and editing, M.J.F., K.R., and J.C.

## Methods

### LEAD CONTACT AND MATERIALS AVAILABILITY

Further information and requests for resources and reagents should be directed to and will be fulfilled by the Lead Contact, Michael Federle. This study did not generate new unique reagents.

### EXPERIMENTAL MODEL AND SUBJECT DETAILS

#### Bacterial strains

*S. pyogenes* (Group A Streptococcus, GAS) strain NZ131 was used as the parental strain for genetic mutants (constructed as described below). Other GAS strains tested included HSC5, 5448, and MGAS315. GAS were routinely grown without shaking at 37° C in Todd-Hewitt 0.2% yeast extract (THY) broth or on agar plates, or in a chemically defined medium (CDM) containing 1% (wt/vol) glucose.^47^ To induce Rgg2/3 activation, 100 nM of synthetic SHP pheromone or inactive reverse SHP (revSHP) was added to *S. pyogenes* cultures at OD_600_~0.1. When appropriate, antibiotics were added at the following concentrations: chloramphenicol (Cm), 3 μg/ml; erythromycin (Erm), 0.5 μg/ml; kanamycin (Kan), 150 μg/ml; spectinomycin (Sp), 100 μg/ml. Starter cultures were utilized to minimize differences in lag phase and were prepared as follows: GAS strains of interest were streaked on THY plates containing the appropriate antibiotics. Single clones were isolated and inoculated into THY containing the appropriate antibiotics. After incubation overnight at 37° C, the cultures were diluted 1:20 into CDM and grown at 37° C to mid-exponential phase (OD_600_ 0.5-0.8). Lastly, glycerol was added to a final concentration of 20%, and single-use aliquots were stored at −80° C.

#### Construction of mutant strains and complementation plasmids

All bacterial strains used in this study are described in Table 1. Primers used to construct plasmids for gene deletion or complementation are listed in Table 2. All cloning was done using laboratory *E. coli* cloning strains such as NEB-5a (New England Biolabs) or BH10c,^48^ with antibiotics added at the following concentrations: Cm, 10 μg/ml; Erm, 500 μg/ml; Sp, 100 μg/ml. To construct gene deletions, sequences flanking the gene of interest were amplified by PCR and ligated into a temperature-sensitive plasmid, pFED760, by restriction enzyme digest or Gibson assembly. When necessary, antibiotic resistance markers for kanamycin (*aphA3)* or chloramphenicol (*cat)* were cloned between the upstream and downstream flanking regions to generate plasmids for selective allelic replacement. Deletion vectors were electroporated into NZ131 and a two-step temperature dependent selection process was used to isolate mutants of interest. Briefly, cells containing each deletion construct were grown at the permissive temperature, then shifted to 37° C and plated on the appropriate antibiotic to select for bacteria in which the plasmid had integrated at one of the flanking regions. After confirmation of plasmid integration by PCR, cells were grown for ~50 generations at the permissive temperature to allow the plasmid to recombine out, and loss of antibiotic resistance was used to identify the desired mutants. Genotypes were confirmed by PCR. To construct complementation plasmids for BGC mutants, the genomic region encompassing *spy49_0450-0457* and its native promoter, were amplified by PCR and cloned into a multi-copy vector (pLZ12-Sp). For complementation of *spy49_0459*, the gene was amplified by PCR and cloned downstream of sequence containing ~70 bp of the *shp3* promoter in an integrating shuttle vector (p7INT/pJC420).

**Table 1.** Strains and plasmids used in this study.

| Strain | Description | Reference |
| --- | --- | --- |
| $\Delta covR$ | NZ131 $\Delta covR$; unmarked | <sup>36</sup> |
| 5448 | Wild-type M1 isolate of the M1T1 lineage | <sup>37</sup> |
| HSC5 | Wild-type M14 isolate | <sup>38, 39</sup> |
| JCC131 | NZ131 $\Delta rgg3::cat$; $cm^R$ | <sup>3</sup> |
| JCC137 | NZ131 $\Delta rgg2$; unmarked | <sup>3</sup> |
| JCC155 | NZ131 $\Delta spy49\_0450-0456$; unmarked | This study |
| JCC173 | NZ131 $\Delta spyCEP::aphA3$; $kan^R$ | This study |
| JCC303 | NZ131 $\Delta spy49\_0450-0460$; unmarked | This study |
| JCC304 | NZ131 $\Delta spy49\_0459$; unmarked | This study |
| JCC306 | NZ131 $\Delta spy49\_0460$; unmarked | This study |
| JCJ134 | NZ131 $\Delta hasAB::cat$; $cm^R$ | This study |
| JCJ173 | NZ131 $\Delta stcA$; unmarked | <sup>4</sup> |
| MGAS315 | Wild-type M3 isolate | <sup>40, 41</sup> |
| NZ131 | Wild-type M49 isolate | <sup>42</sup> |
| RVW113 | NZ131 $\Delta emm::aphA3$; $kan^R$ | This study |
| RVW179 | NZ131 $\Delta slo_{1-260}::aphA3$; $kan^R$ | This study |
| Plasmid | Description | Reference |
| p7INT | Shuttle-suicide vector that integrates at a streptococcal prophage tmRNA site; for complementation in single copy; $erm^R$ | <sup>43, 44</sup> |
| pFED760 | Shuttle vector pGh9-ISS1 deleted for ISS1 element; used for constructing all deletion mutants; temperature-sensitive replication origin; $erm^R$ | <sup>45</sup> |
| pJC202 | To construct unmarked deletion of *spy49_0450-0456*; in pFED760; $erm^R$ | This study |
| pJC233 | Complementation of *spy49_0450-0457* in multi-copy; in pLZ12-Sp; $sp^R$. Nicknamed *p0450-57* in text. | This study |
| pJC258 | To replace *spyCEP* with the *aphA3* kanamycin resistance marker; in pFED760; $kan^R$ $erm^R$ | <sup>36</sup> |
| pJC411 | To construct unmarked deletion of *spy49_0450-0460*; in pFED760; $erm^R$ | This study |
| pJC412 | To construct unmarked deletion of *spy49_0459*; in pFED760; $erm^R$ | This study |
| pJC414 | To construct unmarked deletion of *spy49_0460*; in pFED760; $erm^R$ | This study |
| pJC420 | p7INT-based vector containing 74 bp of *shp3* promoter region; $erm^R$ | This study |
| pJC469 | Complementation of *spy49_0459* in single copy; in pJC420; $erm^R$. Nicknamed *p0459* in text. | This study |
| pJJ148 | To replace *hasAB* with the *cat* chloramphenicol resistance cassette; in pFED760; $cm^R$ $erm^R$ | This study |
| pLZ12-Sp | Shuttle vector for complementation in multi-copy; $sp^R$ | <sup>46</sup> |
| pRVW48 | To replace the *emm49* region with *aphA3* kanamycin resistance marker; in pFED760; $kan^R$ $erm^R$ | This study |
| pRVW82 | To replace the N-terminal 260 amino acids of *slo* with the *aphA3* kanamycin resistance marker | This study |

cm = chloramphenicol; erm = erythromycin; kan = kanamycin; sp = spectinomycin

**Table 2.**
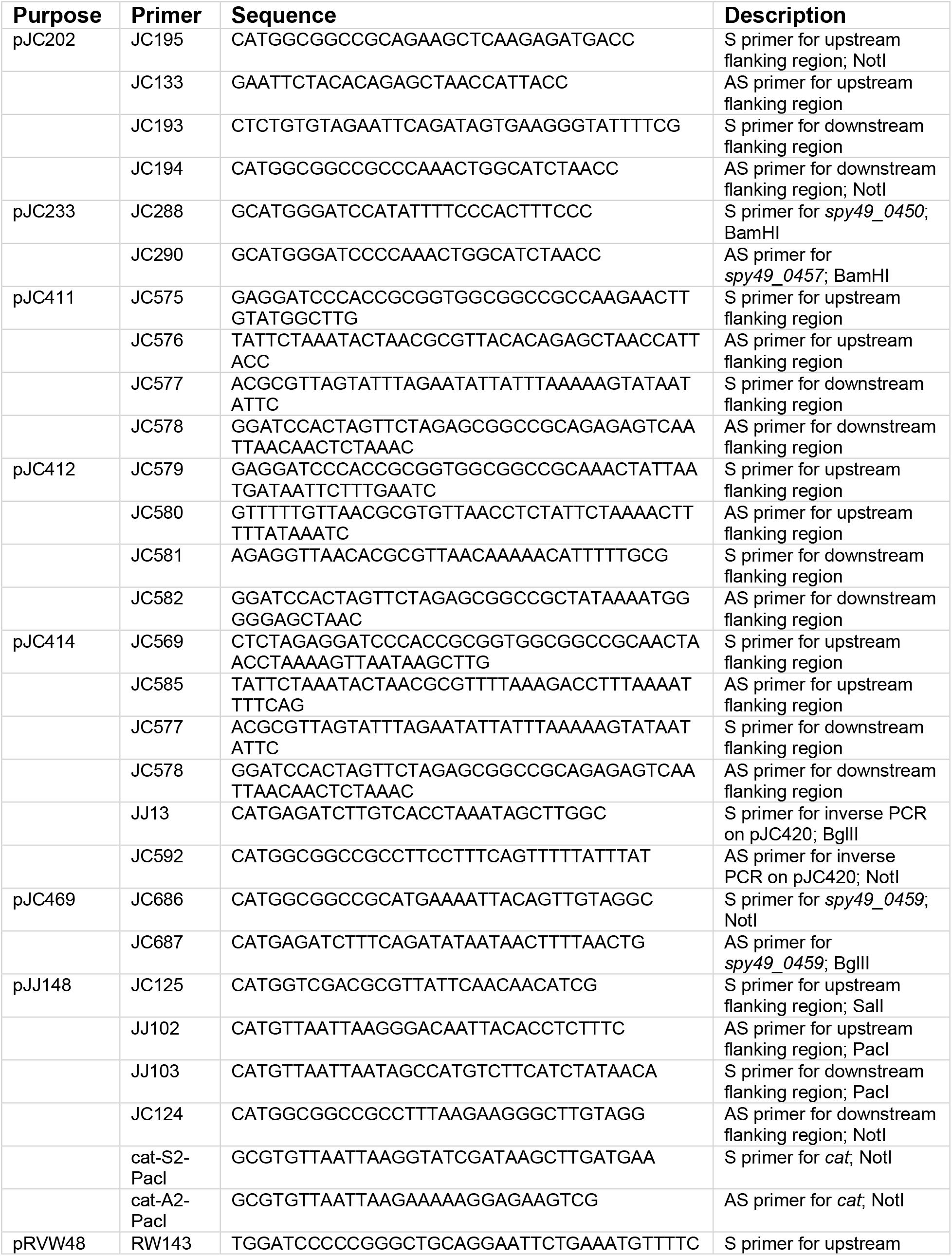
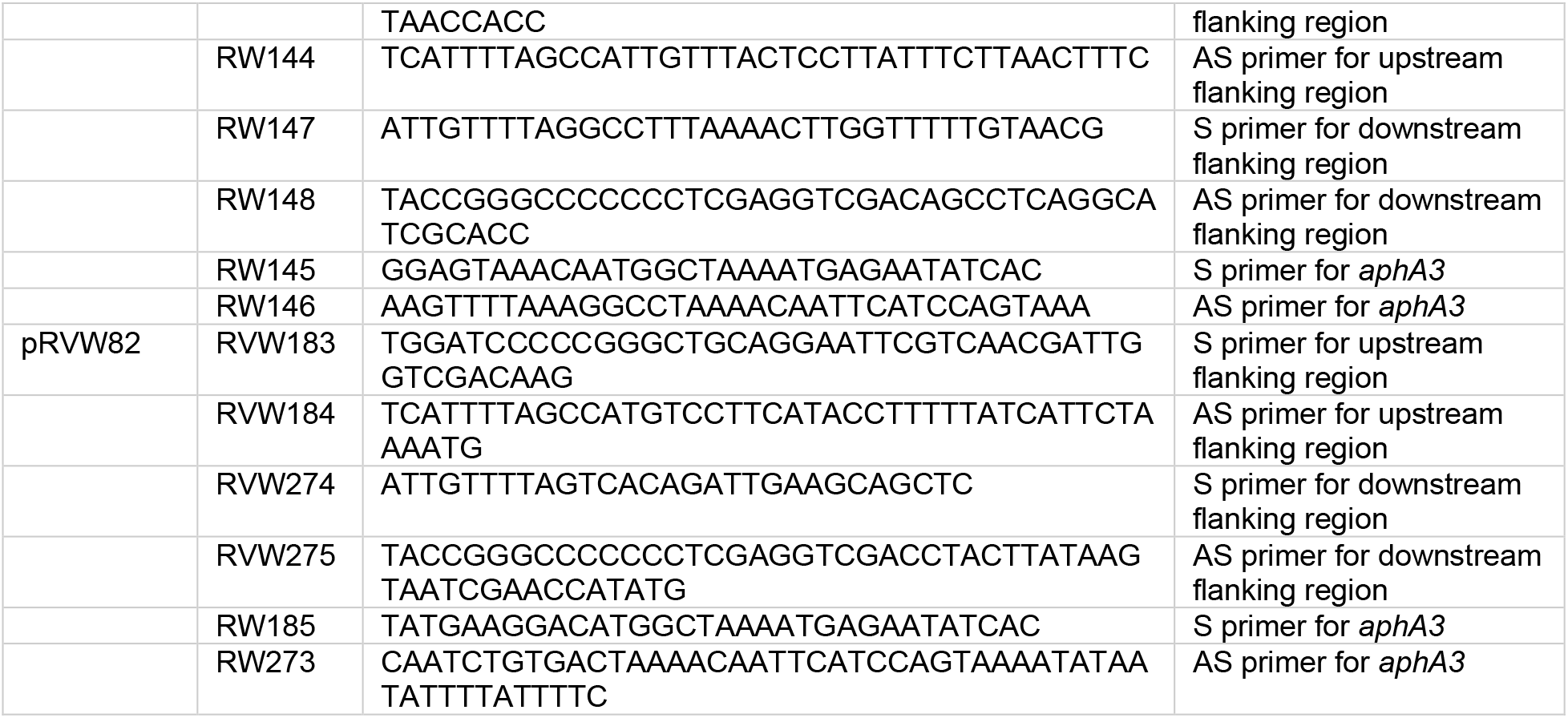
Primers used in this study.

#### Cell lines

NF-κB reporter macrophages (RAW-Blue™ InvivoGen) were cultured in DMEM (Gibco) supplemented with 10% FBS (Gemini, BenchMark), penicillin/streptomycin (Corning) and zeocin (InvivoGen) and RAW264.7 macrophages were maintained in RPMI 1640 (Corning) supplemented with 10% FBS and P/S. All cell lines were maintained and passaged at 37° C + 5% CO_2_.

#### Primary cell culture

BMDM and BMDC were generated from bone marrow precursor cells extracted from femurs and tibiae of 6-8 week-old male or female C57BL/6 mice (Charles River Laboratories).

### METHOD DETAILS

#### Synthetic pheromone peptides

Synthetic peptides of 95% purity were purchased from NeoPeptide (Cambridge, MA), reconstituted as 1 mM stocks in dimethyl sulfoxide (DMSO), and stored at −80° C. Dilutions for working stocks were made in DMSO and stored at −20° C. The SHP3-C8 peptide (SHP) sequence is DIIIIVGG, and the reverse peptide (rev-SHP) sequence is GGVIIIID.

#### *In vitro* infections

2.5 × 10^5^ macrophages were seeded into 24-well tissue culture treated plates (Corning) in 500 μL of medium supplemented with penicillin/streptomycin the day before infection. The following day, medium was aspirated and replaced with medium without antibiotics. GAS was grown from starter cultures in CDM to OD_600_= 0.5 (~2-3 hours), then normalized based on OD, washed in PBS, and added to macrophages. Unless otherwise noted, macrophages were inoculated with GAS at a multiplicity of infection of 10:1. Cells were immediately centrifuged at 200 × g for 5 minutes to equilibrate infection and then incubated at 37° C with 5% CO_2_. After 30 minutes, extracellular bacteria were killed by replacing with medium containing 100 μg/ml gentamicin (Gibco). Where applicable, cells were treated with the following TLR agonists from InvivoGen: Lipopolysaccharide (LPS, 100 ng/ml), Heat-Killed *Listeria monocytogenes* (HKLM, 1 × 10^8^ cells), or CpG oligodeoxynucleotides (ODN1826, 1 μM). For experiments with attenuated GAS, bacteria were killed with heat (65C), UV (short wave), or 4% PFA for 60 minutes. For experiments using GAS supernatants, cells were grown to OD_600_~0.5 and pelleted, and the supernatants were subsequently filtered (0.2 μm) and normalized based on OD. For trans-well experiments, 0.4 μm pores (Corning) were used.

For assays where phagocytosis was blocked, macrophages were pre-treated with 0.5 μg/ml Cytochalasin D (CytD) for 1 hour. After inoculating with GAS for 30 minutes, cells were treated with gentamicin (400 μg/ml) for 30 minutes to kill remaining GAS. After washing three times with PBS to remove the remaining GAS, gentamicin, and CytD, cells were then stimulated with either LPS (100 ng/ml) or GAS (MOI 10).

#### Cell viability assay

Cytotoxicity of macrophages after 8 hours of infection was quantified by measuring the release of the enzyme lactate dehydrogenase in cell supernatants using the CytoTox 96 Non-Radioactive Cytotoxicity Assay (Promega) according to the manufacturer’s instructions. Percentages were determined by comparing samples with a 100% lysis control.

#### NFκB activation assay

RAW Blue macrophages (InvivoGen) with chromosomally integrated NFκB inducible secreted embryonic alkaline phosphatase (SEAP) reporter construct were used to measure activation of NFκB. Cells were maintained in DMEM (Gibco) supplemented with 10% heat inactivated FBS (Gemini) and penicillin/streptomycin (Corning) and 200 μg/ml zeocin (InvivoGen) in tissue culture treated T25 flasks (Greiner Bio-One). Cells were grown to 60-70% confluency, washed with PBS, and then passaged using 0.05% trypsin/0.53 mM EDTA (Corning) to dissociate them from the flasks. Reporter macrophages were counted and seeded the day prior to the experiment in flat bottom 96-well tissue culture treated plates (Corning) at 5 × 10^4^ cells/well. The following day, GAS was added to the wells at a multiplicity of infection of 10:1 and cells were centrifuged for 5 minutes to equilibrate infection. After a 30-minute incubation at 37° C, the medium was removed and replaced with medium containing 100 μg/ml gentamicin. After 18 h, RAW Blue cell-free supernatants were collected and 50 μl was mixed with 150 μl of the QUANTI-Blue substrate (InvivoGen) and incubated at 37° C for 45 minutes in a flat bottom 96-well plate. Color change was quantified by measuring absorbance at 625 nm using a Synergy HTX microplate reader (BioTek).

#### Generation of Bone Marrow-derived Macrophages and Dendritic Cells

Primary murine bone marrow-derived macrophages (BMDMs) and dendritic cells (BMDCs) were differentiated from bone marrow cells collected from the femurs and tibiae of six to eight-week-old male C57BL/6 (Charles River Laboratories). For macrophage differentiation, bone marrow cells were plated in 100 × 15 mm petri dishes in 10 mL DMEM supplemented with 10% FBS, 1 mM HEPES buffer (Gibco), 1x Penicillin/Streptomycin (Corning) and 10 ng/ml MCSF (BioLegend) and incubated at 37° C + 5% CO_2_. On day 3, medium was removed and cells were re-supplemented with 10 mL fresh medium. On day 7, the cells were washed gently with PBS and incubated with trypsin for 5 minutes at 37° C. Cells were then removed by gentle pipetting, counted, and plated at 2.5 × 10^5^ cells per well on 24 well plates or 5 × 10^4^ cells per well on 96 well plates overnight and infected on day 8.

For dendritic cell differentiation, red blood cells were lysed using RBC Lysis Buffer (Tonbo) and following a centrifugation step, cells were counted and plated on 6 well plates in RPMI + 5% FBS + 20 ng/ml GM-CSF (BioLegend). Cells were maintained at 37° C 5% CO_2_ and supplemented on day 3, 6, and 8. On day 10 cells were supplemented with medium containing 12.5 ng/ml GM-CSF. Cells were collected on day 11 and infected on day 12.

The purity of BMDMs and BMDCs was measured by cell surface staining and flow cytometry. 2.5 × 10^5^ cells were transferred to an Eppendorf tube and resuspended in FACS buffer (PBS + 1% BSA). Prior to staining, cells were treated with anti-CD16/CD32 (BioLegend) for 15 minutes to block Fc receptors. Cells were then stained for 30 minutes on ice with anti-CD11b-APC/Fire 750 and anti-F4/80-PE for macrophages or anti-CD11c-Alexa Fluor 647 for dendritic cells (BioLegend). Samples were washed twice and then run on Cytoflex (Beckman Coulter). Isolated cells were determined to be >90% pure.

#### Quantification of cytokines

Cell-free culture supernatants were harvested at 8 hours post infection and frozen at −20° C. The levels of murine IL-6 and TNF-α and human IL-6 were measured using ELISA MAX™ kits (BioLegend) according to the manufacturer’s instructions. For murine IFNβ the DuoSet ELISA kit (R&D systems) was used.

#### Measurement of GAS adherence to macrophages

Adherence to RAW264.7 cells was measured after 30 minutes of infection. Cells were gently washed three times with 500 μL PBS, followed by incubation with 100 μL trypsin for 5 minutes at 37° C and then addition of 400 μL 0.1% saponin in sterile water for 1-2 minutes until cells appeared lysed under the microscope. Serial dilutions were plated on THY agar plates and CFUs were enumerated after overnight incubation at 37° C with 5% CO_2_.

### QUANTIFICATION AND STATISTICAL ANALYSIS

Data were statistically analyzed using GraphPad Prism v.7.0b. Data were analyzed using ordinary one-way ANOVA followed by Tukey’s multiple comparisons test or two-way ANOVA followed by Sidak’s multiple comparisons test and deemed significant at *p < 0.05; **p < 0.005; ***p < 0.001; ****p < 0.0001. ns is used to denote non-significant comparisons.

## DATA AND CODE AVAILABILITY

This study did not generate/analyze datasets/code.

